# Structure of the human volume regulated anion channel

**DOI:** 10.1101/323584

**Authors:** J. M. Kefauver, K. Saotome, A. E. Dubin, J. Pallesen, C.A. Cottrell, S.M. Cahalan, Z. Qiu, G. Hong, C.S. Crowley, T. Whitwam, W.H. Lee, A.B. Ward, A. Patapoutian

**Affiliations:** Howard Hughes Medical Institute, Department of Neuroscience, The Scripps Research Institute, La Jolla, CA 92037, USA.; Department of Integrative Structural and Computational Biology, The Scripps Research Institute, La Jolla, California 92037USA.; Genomics Institute of the Novartis Research Foundation, San Diego, California 92121USA.

## Abstract

SWELL1 (LRRC8A) is the only essential subunit of the Volume Regulated Anion Channel (VRAC), which regulates cellular volume homeostasis and is activated by hypotonic solutions. SWELL1, together with four other LRRC8 family members, forms a vastly heterogeneous cohort of VRAC channels with different properties; however, SWELL1 alone is also functional. Here, we report a high-resolution cryo-electron microscopy structure of full-length human homo-hexameric SWELL1. The structure reveals a trimer of dimers assembly with symmetry mismatch between the pore-forming domain and the cytosolic leucine-rich repeat (LRR) domains. Importantly, mutational analysis demonstrates that a charged residue at the narrowest constriction of the homomeric channel is an important pore determinant of heteromeric VRAC. This structure provides a scaffold for further dissecting the heterogeneity and mechanism of activation of VRAC.

## Main Text

VRAC is a ubiquitously expressed mammalian anion channel implicated in diverse physiological processes including volume regulation, cell proliferation, release of excitatory amino acids, and apoptosis (*1*–*3*). It is suggested to play a role in a variety of human diseases including stroke, diabetes, and cancer (*3*–*5*). A causative link has been established between a chromosomal translocation in the Swell1 (Lrrc8a) gene and a human B cell deficiency disease, agammaglobulinemia (*6*).

Previous studies have shown that SWELL1 is required for VRAC activity, and that the presence of other LRRC8 subunits dictates functional characteristics of VRAC, including pore properties (*7*–*9*). While SWELL1 and at least one other LRRC8 subunit are required for canonical whole-cell VRAC currents, purified homomers of SWELL1 reconstituted in lipid bilayers are activated by osmotic stimuli and blocked by VRAC antagonist, DCPIB (*9*). Interestingly, CRISPR-engineered HeLa cells lacking all LRRC8 subunits (LRRC8^−/−^ HeLa cells) exhibited very small but significant DCPIB-sensitive hypotonicity-induced currents after SWELL1 overexpression (Fig. S1), supporting previous bilayer results. Since the number and composition of functional native oligomeric assemblies remains unknown, we decided to first elucidate the structure of SWELL1 homomers. To produce homomeric SWELL1, human SWELL1-FLAG was recombinantly expressed in LRRC8(B,C,D,E)^−/−^ HEK293F suspension cells, then detergent-solubilized and purified for structure determination by cryo-EM (Fig. S2). Image analysis and reconstruction yielded a ~4 Å resolution map that was used to build a molecular model of SWELL1 (Fig. S3-S4, Table S1).

SWELL1 is organized as a hexameric trimer of dimers with a four-layer domain architecture and an overall jellyfish-like shape (**Fig. 1A**). The transmembrane (TM) and extracellular domains (ECDs) surround the central pore axis, and share a previously unappreciated structural homology with the connexin(*10*) and innexin(*11*) gap junction channels (Fig. S5A-D). The ECD is composed of two extracellular loops (ECL1 and ECL2) that are stabilized by three disulfide bonds (**Fig. 1B-C** and fig. S5E-F). Each subunit contains four TM helices (TM1-4). TM1 lies closest to the central pore axis and is tethered to a short N-terminal coil (NTC) that is parallel to the membrane. In the cytosol, the intracellular linker domains (ILD) create a tightly packed network of helices connecting the channel pore to the LRR domains. Each ILD is composed of four helices from the TM2-TM3 cytoplasmic loop (LH1-4), and five helices from the TM4-LRR linker (LH5-9) (**Fig. 1C**). Each protomer terminates in 15–16 LRRs which form a prototypical solenoid LRR fold (**Fig. 1B-C**). LRRs on three sets of neighboring protomers then dimerize into three pairs, which interact to form a Celtic knot-like assembly (**Fig. 1A**).

**Fig. 1.**
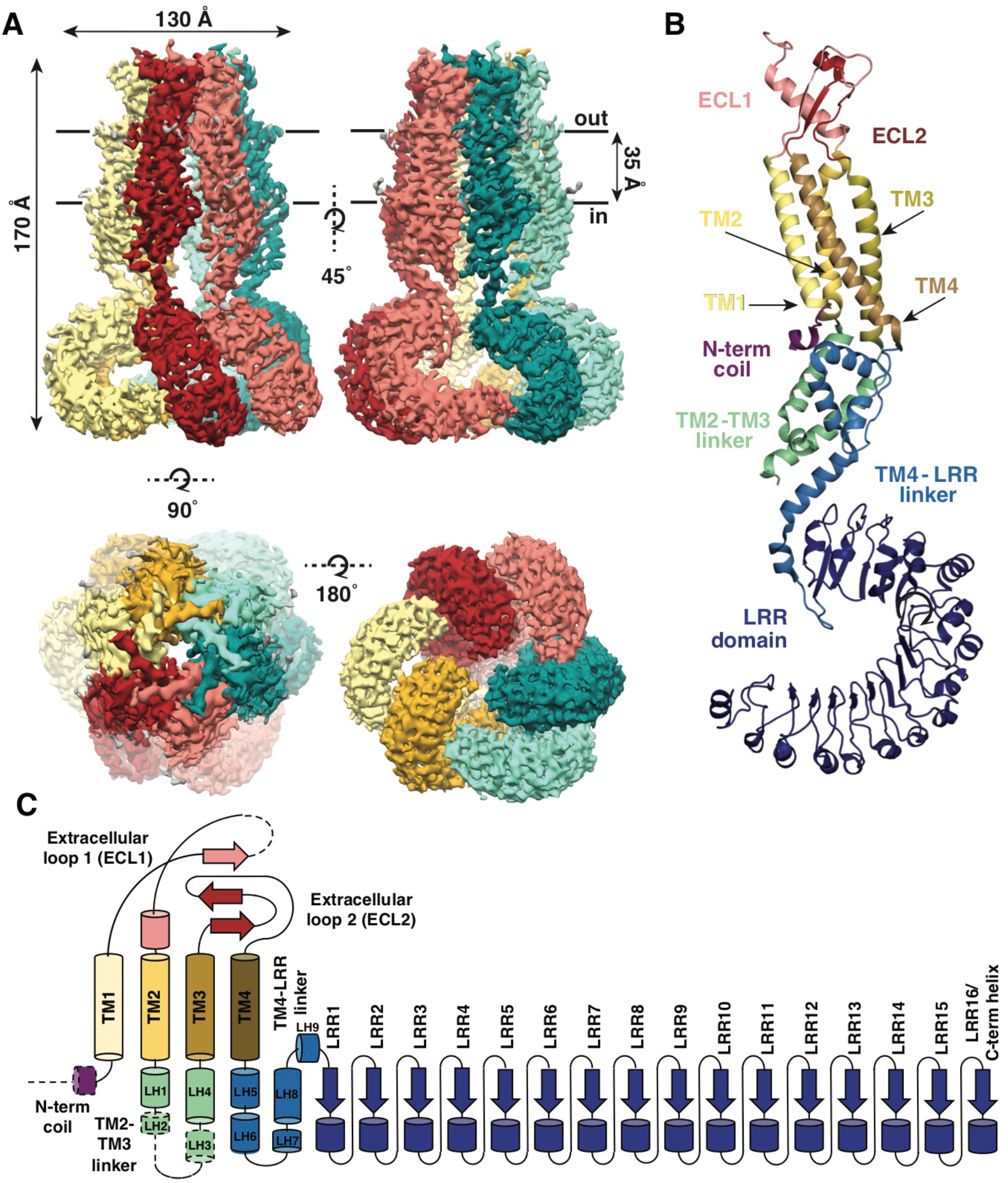
Overall architecture of homomeric SWELL1. (**A**) Cryo-EM reconstruction of SWELL1 homohexamer viewed from the membrane plane highlighting a dimer pair (top left, red and pink subunits) and an interface between dimers (top right, pink and green subunits), from the extracellular side (bottom left), and from the cytosolic side (bottom right). (**B**) Detailed view of SWELL 1 “inner” protomer. (**C**) Topology diagram denoting secondary structural elements. Dashed lines indicate unresolved regions on both protomers in a dimer pair, while dashed shape borders indicate regions that are only resolved on one protomer.

Perhaps the most striking architectural feature of VRAC is the symmetry mismatch between the cytosolic LRR domains and the pore-forming domains of the channel, despite its homo-hexameric assembly (**Fig. 2**). The ECDs, TMs, and ILDs all share the same 6-fold symmetric arrangement (**Fig. 2B**); however, in the cytosol, LRR domains dimerize in a parallel fashion with each LRR at either a 10° or −20° offset relative to the rest of its protomer, producing a 3-fold symmetric trimer of dimers (**Fig. 2C**). The nonequivalence between identical subunits arises from a hinge around the conserved residue L402 in a helix of the TM4-LRR linker (**Fig. 2D** and fig. S6). This hinge allows the LRR domains to shift as rigid bodies, producing sufficient flexibility for them to interface at their edges via several charged residues (**Fig. 2D**, **3A**). As a result, the helical C-termini of the two subunits in a dimer pair make two different sets of interactions with the neighboring LRR (**Fig. 3B**). Focused 3D classification of the LRR domains revealed several arrangements of LRRs suggesting that flexibility of the LRR domains may play a functional role in channel gating (Fig. S7). Interestingly, the outer LRR subunit in the dimer exhibits helical density in the C-terminal half of the TM2-TM3 linker that rests on top of the outer protomer’s LRR domain, adding an additional layer of intricacy to the network of cytosolic interactions (Fig. S5A-B). Symmetry mismatch is also observed in the homotetrameric AMPA receptor GluA2, which similarly forms local dimers in different domain layers (*12*). Furthermore, the dimer-of-dimers topology of homotetrameric AMPA-subtype ionotropic glutamate receptors (iGluRs) defines the subunit organization of di- and tri-heteromeric NMDA-subtype iGluR structures (*13*–*15*). By analogy, we speculate that the trimer-of-dimers assembly of SWELL1 is recapitulated in, and influences the composition of, heteromeric VRACs.

**Fig. 2.**
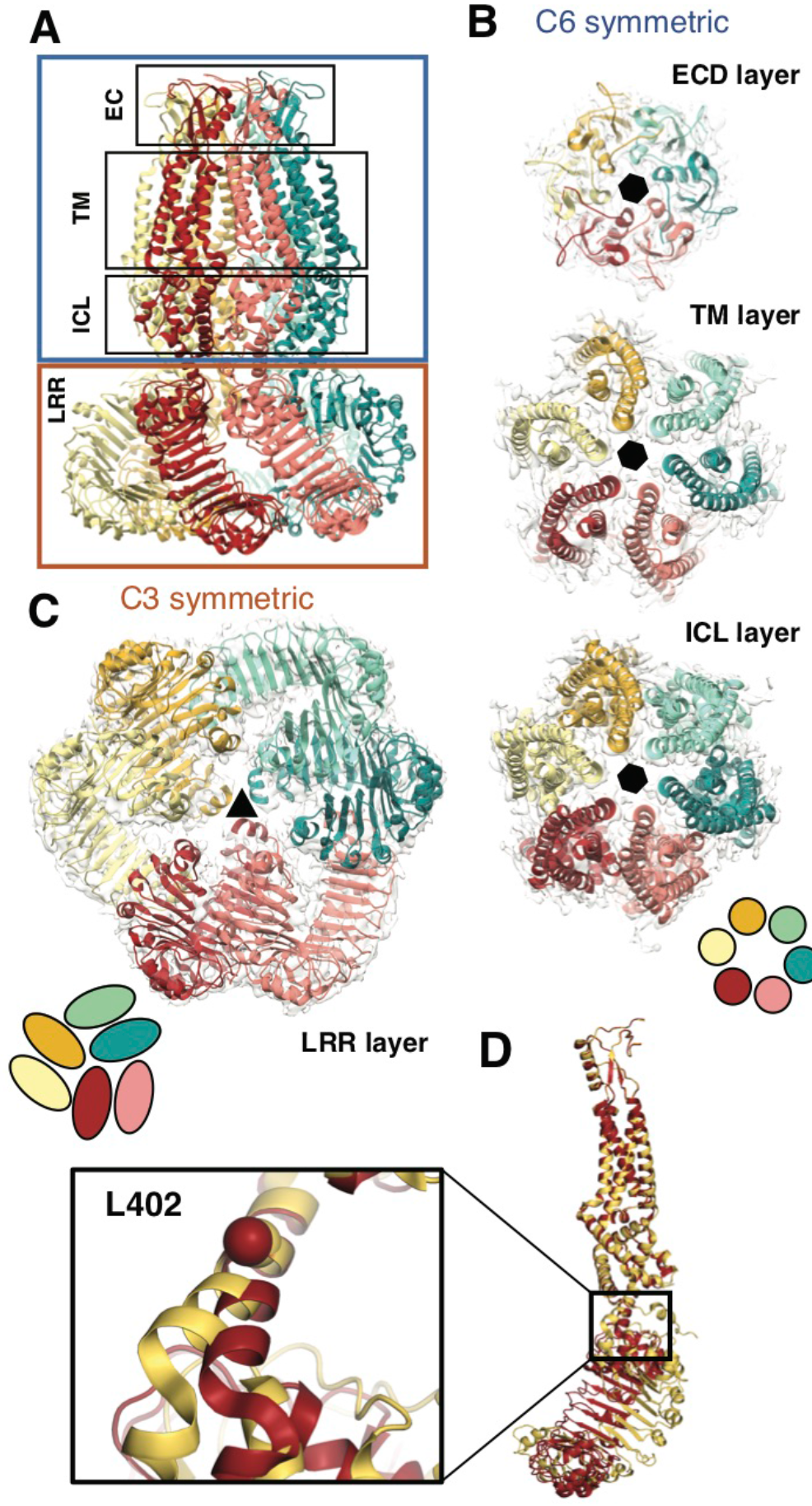
Subunit arrangement exhibits symmetry mismatch. (**A**) SWELL1 model viewed from the membrane plane with domain layers viewed perpendicular to the symmetry axis. (**B-C**) Domain layers viewed from the top of the channel grouped according to shared symmetry with 5 simple schematic to demonstrate subunit arrangement. (**B**) From left to right: extracellular domain layer (EC), transmembrane domain layer (TM), and intracellular linker domain layer (ICL) all share the same 6-fold rotation symmetry axis (black hexagon). (**C**) The LRR domain layer has 3fold rotational symmetry (black triangle), resulting from parallel pairing of three sets of LRR domains. (**D**) Asymmetry in LRR pairing arises from a hinge at L402 that allows rotation of the 10 LRR domain as a rigid body in a dimer pair.

**Fig. 3.**
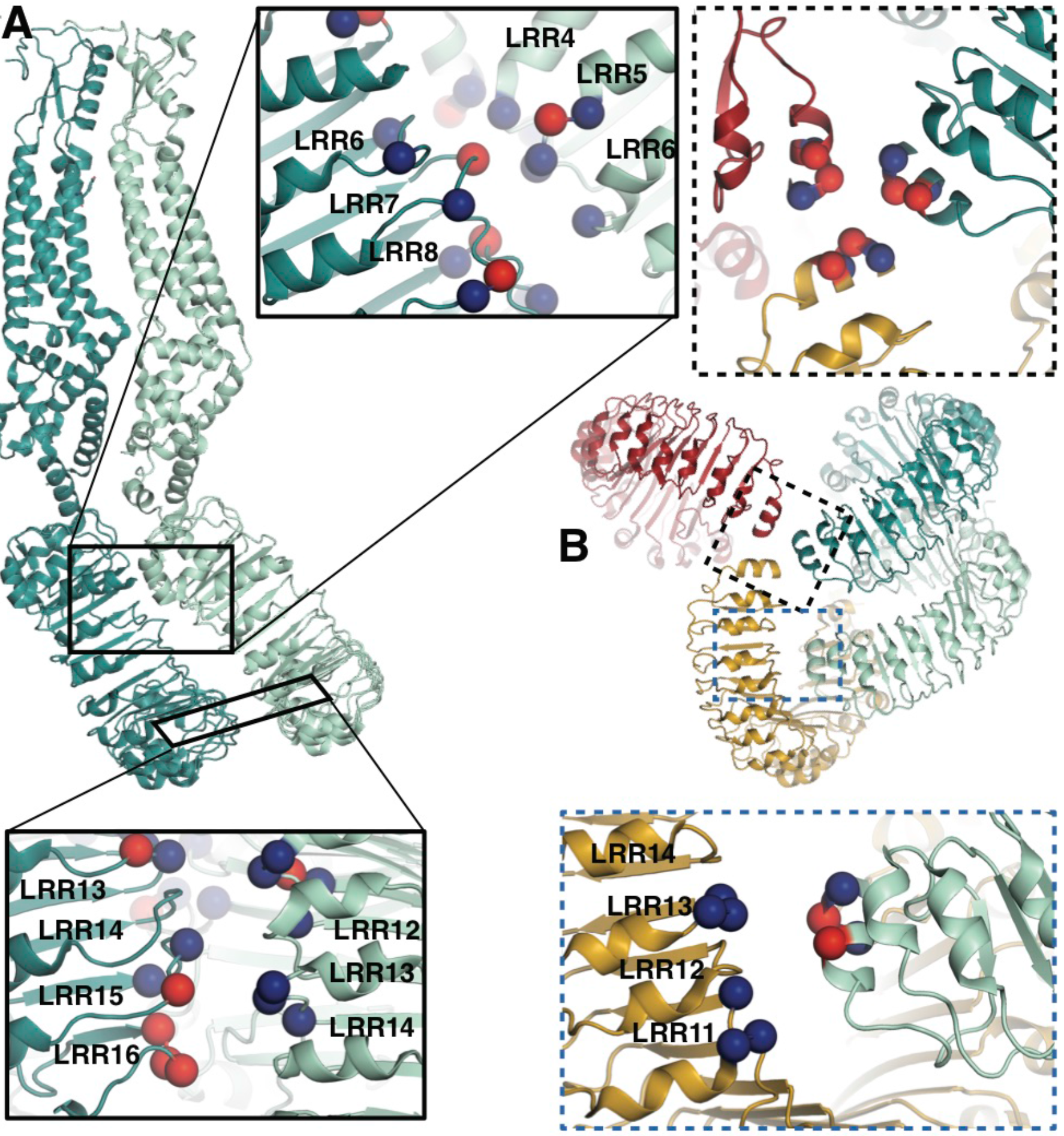
LRRs interact via charged residues at dimer interfaces and C-termini. (**A**) One dimer of SWELL 1 subunits. Charged residues both of opposite and similar charges face each other in the interface between the two LRR domains (insets, top middle and bottom left; blue spheres are 5 positively charged residues (Arg, Lys, and His), red spheres are negatively charged residues (Asp and Glu)). (**B**) C-termini of the two protomers in a dimer interact with regions of the neighboring LRR domain. Two of three “outer” subunits are removed for clarity. “Inner” subunits may be able to coordinate with one another via a triad of charged residues (E800) at their C-termini (inset, dashed border, top right), while the C-termini of the “outer subunit” may interact with the edge of 10 the neighboring outer subunit via charged residues R688 on LRR12 and/or R732 on LRR13 (inset, dashed blue border, bottom right).

Unlike other ion channels, there is little domain swapping between the subunits of the pore-forming domains of the SWELL1 channel. The individual helical bundles are loosely packed with one another and lined with hydrophobic residues. Within the membrane, the most prominent subunit interface is at a pair of hydrophobic residues (F41 and Y127). To investigate whether this region in VRAC is involved in stabilizing the channel, we mutated Y127 to cysteine. Overexpression of SWELL1-Y127C together with LRRC8C produced a constitutive DCPIB-sensitive conductance (Fig. S8A-C). Also, in the TM region near the extracellular face, annular densities occupy the inter-subunit space between each protomer (Fig. S8D). These likely correspond to lipids or cholesterol carried over from purification, or well-coordinated detergent molecules. Similar densities are observed in the inter-subunit space in innexin-6 and have been proposed to have a stabilizing role in the conformation of the helix bundles(*11*). This region and its interaction with hydrophobic membrane components like lipid or cholesterol may be important for channel assembly or lipid signaling.

At the upper faces of the extracellular domains, on mostly flexible loops, resides the three residue KYD motif involved in voltage-dependent inactivation and selectivity(*16*); interestingly, KYD extends laterally towards the neighboring subunit (Fig. S9A), suggesting that subunit interactions in this region contribute to these channel properties.

The ECDs, TMs, and ILDs of all six subunits contribute to the ion-conducting pore (**Fig. 4A-B**). Below that, windows of 35 Å by 40 Å between LRR dimer pairs are sufficiently large to allow ions and osmolytes to freely pass. In the extracellular domain, a ring of arginines (R103) forms the narrowest constriction in the channel structure (**Fig 4A-C**). We hypothesized that these arginines, only conserved between SWELL1 and the LRRC8B subunit (R99) (Fig. S6), might directly interact with permeant anions. To test this hypothesis, we mutated positively-charged R103 to phenylalanine, and determined whether ion selectivity was altered in SWELL1-R103F + LRRC8C heteromeric channels heterologously expressed in HeLa (ABCDE)^−/−^ cells. We determined the reversal potential (V_rev_) for hypotonicity-induced Cl^−^ currents mediated by SWELL1-R103F + LRRC8C channels. The V_rev_ of currents mediated by SWELL1-R103F + LRRC8C was significantly reduced compared to wildtype channels, indicating that the channels are less selective for Cl^−^ (**Fig 4D**) (*17–19*). Furthermore, extracellular ATP at concentrations that block ~75% of wildtype VRAC currents was ineffective on channels containing R103F (**Fig. 4E** and fig S9B). Therefore, R103 is a critical residue within SWELL1 that impacts ion selectivity as well as pore block of heteromeric VRAC channels.

**Fig. 4.**
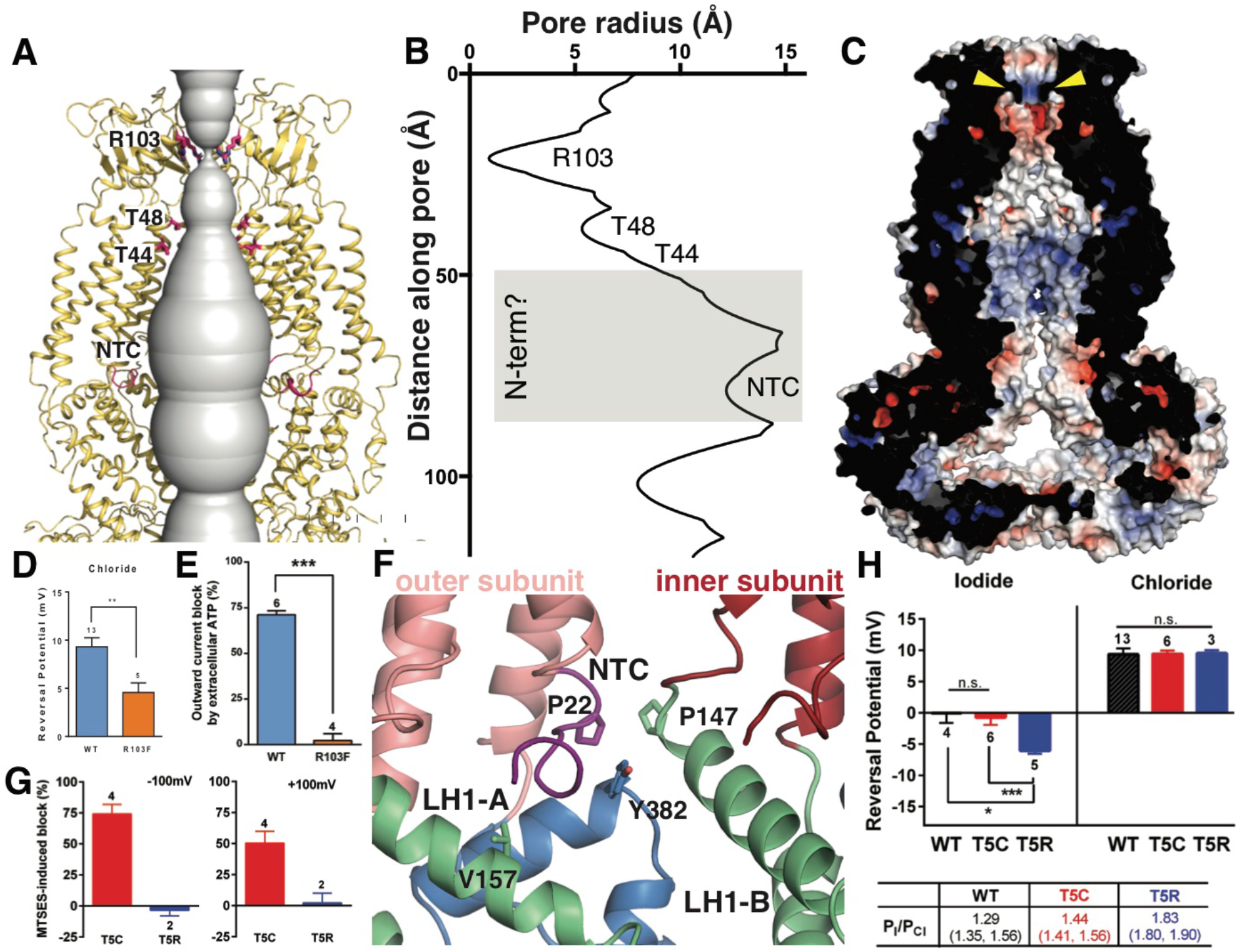
Ion pore structure and electrophysiological characterization of residues R103 and T5. Cartoon model of the SWELL1 pore, with two subunits removed for clarity. A surface representation of the radial distance between the protein surface and the pore axis is shown in grey. Pore-facing residues R103, T48 and T44, and N-terminal coil (NTC) are labeled in pink. (**B**) Graph of van der Waals radii of the pore, plotted against distance along the pore axis. Locations of residues R103, T48, T44, and NTC are labeled along 2D plot. Grey box covers potential area the N-terminus might occupy. (**C**) Electrostatic surface potential of channel pore, viewed by vertical cross-section. Narrow constriction on the extracellular side of the channel is formed by a ring of R103 residues (yellow arrows). Calculated using APBS implemented by Pymol2.0 with potentials ranging from -10 kT (red) to +10 kT (blue). (**D**) Cl- selectivity is reduced in R103F (V_rev_:+4.6 ± 1.0 mV (n=5)) relative to WT (V_rev_: +9.4 ± 0.9 mV (n=13), P < 0.005; Student’s *t*-test). Mean ± S.E.M. for the number of cells indicated. (**E**) R103F-containing channels are insensitive to block by externally applied Na_2_ATP. Whole cell currents induced by hypotonic solution (230 mOsm/kg) were measured at +100 mV before and during 2 mM ATP exposure and leak subtracted. For cells expressing wildtype SWELL 1-containing channels outward currents were rapidly reduced by ATP block, while essentially no ATP block was observed in cells expressing R103F channels. (**F**) Detailed view of coordination of NTC (purple). The NTC makes intrasubunit contacts with V157 on LH1 and a conserved Y382 at the kink between LH6 and LH7 of the TM4-LRR linker. Additionally, P22 of the NTC makes an intersubunit contact with a conserved P147 at the kink between TM2 and the TM2-TM3 linker of the neighboring subunit. (**G**) The polar MTS reagent MTSES applied extracellularly blocks SWELL1-T5C containing channels but has no effect on T5R containing channels that are not modifiable by MTS reagents. Bar graphs show maximum percent block (mean ± S.E.M. for the number of cells indicated) during exposure to MTSES at either -100mV (left) or +100mV (right). The percent block by 3.33 mM MTSES of T5C-mediated currents is 74.2 +/- 7.7% and 50.4 +/- 9.5% (n=4) at -100 and +100 mV, respectively (mean +/- S.E.M.). (**H**) Relative permeability PI/PCI is enhanced by the T5R mutation. Reversal potentials in iodide and chloride for hypotonic solution (230 mOsm/kg)-induced currents mediated by the indicated SWELL1 construct together with LRRC8C. The calculated PI/PCI is shown in the table (mean with lower and upper 95% confidence intervals). Whole cell experiments were performed on HeLa (ABCDE)^−/−^ cells transfected with SWELL1 (or the indicated point mutation) and LRRC8C-ires-GFP.

Within the pore, constrictions are observed at pore-facing residues T44 and T48 (**Fig 4A-B**). Interestingly, we had previously identified residue T44 via the substituted cysteine accessibility method (SCAM) on heteromeric channels as likely to be at or near the pore (*8*). Near the bottom of the pore cavity, a constriction at the intracellular face of the membrane corresponds to a short N-terminal coil (NTC) sitting parallel to the membrane. The first 14 residues of the N-terminus of the channel are not resolved in the cryo-EM density, presumably due to flexibility. The absence of these residues is conspicuous; in the Cx26 and innexin-6 structures, an N-terminal helix forms a pore funnel structure that is the narrowest constriction in the structures of these channels and is thought to contribute to trafficking, selectivity, and gating(*10*, *11*, *20*, *21*). In our reconstruction, the short portion of the NTC that is resolved is highly coordinated by cytosolic domains and positioned to respond to conformational changes in the cytosolic domains of one protomer, as well as movements of the neighboring protomer (**Fig. 4F**). Due to the similarities in pore structure between VRAC and connexin/innexin (Fig. S5), we conducted functional assays to interrogate the role of the NTC in VRAC. We focused on residue T5 because this residue is involved in stabilizing the pore funnel through a hydrogen bonding network in the Cx26 structure(*10*). We made the mutation T5C to test whether extracellular addition of the negatively-charged, membrane-impermeable thiol-reactive reagent, 2-sulfonatoethyl methanethiosulfonate (MTSES), could alter VRAC activity in HeLa LRRC8(A,B,C,D,E)^−/−^ via cysteine modification. While MTSES has no effect on wildtype heteromeric channels (*8*) or channels containing T5R, whole-cell currents mediated by SWELL1-T5C + LRRC8C are strongly suppressed upon the addition of MTSES, suggesting that T5C is part of a constriction narrow enough to block the pore upon covalent modification by MTSES (**Fig. 4G** and Fig. S9C). We next determined the role of T5 in anion selectivity. Although T5C-containing channels have similar relative permeability to wildtype, T5R-containing channels are significantly more selective to iodide compared to chloride, confirming that this residue is close to or part of the channel pore (**Fig. 4H** and Fig S9D). Thus, the unresolved portion of the N-terminus plays a role in pore constriction in native channels composed of SWELL1 and LRRC8C. Its absence in our structure is likely due to the high flexibility of the region or a peculiarity of the homomeric assembly of the channel.

Here we report the architecture and homo-hexameric assembly of SWELL1 channels. Electrophysiological analyses presented here demonstrate that the homomeric SWELL1 structure retains properties of the more complex heteromers, as mutations based on the structure proved to be relevant for VRAC currents in a cellular context. The structure of SWELL1 also provides hints as to how VRAC gating is regulated. Since decreases in intracellular ionic strength cause activation(*9*), gating would likely be initiated by movement of intracellular domains in response to changes in salt concentration. We speculate that the multitude of charge-mediated interactions in the LRRs endows the SWELL1 structure with ionic-strength sensitivity, and the ILDs couple LRR movement to the transmembrane channel.

## Acknowledgments

We thank H. Turner, W. Anderson, C. Bowman, and T. Nieusma for training in electron microscopy and computational methods. We acknowledge A. Coombs for molecular biology assistance. We thank R. MacKinnon, G. Lander, S. Murthy and members of the Ward and Patapoutian labs for helpful discussions. This work was supported by National Institutes of Health (NIH) National Research Service Award F31 NS093778-3 to J.M.K., a Ray Thomas Edwards Foundation grant to A.B.W., and NIH grant NS083174 to A.P. A.P. is an investigator of Howard Hughes Medical Institute (HHMI).

## Author contributions

JMK, GH, ZQ and CSC developed sample preparation protocols. SMC and ZQ generated CRISPR cell lines. JMK expressed and purified protein, prepared cryo-EM samples, and collected images. JP and JMK processed cryoEM data. KS, JMK, and CAC built and refined atomic models. AED designed and performed the whole cell electrophysiological experiments. GH, TW, and WHL created mutant constructs and provided molecular biology technical assistance. JMK, KS, AED, ABW, and AP analyzed the results and wrote the manuscript, with all authors editing and approving the final manuscript.

## Data and materials availability

The cryo-EM map of human SWELL1 was deposited into the Electron Microscopy Data Bank with accession code XXXX. The atomic model of human SWELL1 was deposited into the Protein Data Bank with PDB ID XXXX.

## Materials and Methods

### CRISPR LRRC8(B,C,D,E) KO cell line

Knock-out of LRRC8 genes in HeLa cell line and HEK293F suspension cell line was completed using CRISPR/Cas9-mediated gene disruption (*22*). LRRC8A, LRRC8B, LRRC8D, and LRRC8E genes were targeted using guideRNA (gRNA) sequences reported by Voss, et al., 2014; the LRRC8C gene was targeted with a gRNA sequence reported by Syeda, et al, 2016 (*9*). Cloning of the gRNAs into PX458-mCherry plasmid was completed as reported in Syeda, et al, 2016 (*9*). Multiple plasmids were transfected simultaneously using either Lipofectamine 2000 or PEI max. After 48-72 hr, fluorescent mCherry positive cells were single-cell sorted into 96-well plates. Successful knock-out was determined by genotyping targeted regions for frameshift mutations and verified by mass spectrometry analysis. For HeLa cells (LRRC8^−/−^ HeLa cells), complete knockout was verified for all five LRRC8 genes. For HEK293F suspension cells, complete knock-out was verified for LRRC8B-E (LRRC8(B,C,D,E)^−/−^ HEK293F cells). One LRRC8A gene locus remained intact in all surviving suspension culture lines.

### Protein expression and purification

Human LRRC8A (Origene #RC208632) was cloned with a C-terminal FLAG-tag (DYKDDDDK) separated by a triple glycine linker (LRRC8A-GGG-FLAG) into a pcDNA3.1/Zeo(-) vector using Gibson cloning. HEK293F *LRRC8(B,C,D,E)-/-* cells were transfected at a cell density of 1.8*10^6 cells/mL with 1mg/L of LRRC8A-GGG-FLAG plasmid DNA combined with 3mg/L cells of PEI max. After 48 hours, cells were pelleted and solubilized in solubilization buffer (20mM Tris pH 8, 150mM NaCl, 1% DMNG, 2mg/mL iodoacetamide, and EDTA-free protease inhibitor cocktail (PIC)) at 4°C with vigorous shaking. The cell lysate was centrifuged at 90*xg* for 30 min at 4°C and the supernatant was collected and combined with 1mL/L cells of FLAG M2 affinity resin for 1 hr batch incubation at 4°C with gentle shaking. Resin was washed in a gravity column with 5CV of solubilization buffer (20mM Tris pH 8, 150mM NaCl, 1% DMNG, 2mg/mL iodoacetamide, and EDTA-free PIC), 5CV of high salt wash buffer (20mM Tris pH 8, 150mM NaCl, 0.05% digitonin, and EDTA-free PIC), and 10CV of wash buffer (20mM Tris pH 8, 150mM NaCl, 0.05% digitonin, and EDTA-free PIC). Protein was eluted using elution buffer (20mM Tris pH 8, 150mM NaCl, 0.05% digitonin, EDTA-free PIC and 3x FLAG peptide. Sample was concentrated and injected onto Shimadzu HPLC and separated using a Superose 6 Increase column equilibrated with running buffer (20mM Tris pH 8, 150mM NaCl, 0.05% digitonin, and EDTA-free PIC). The peak corresponding to LRRC8A homomeric oligomers (~800kDa) was collected and used for cryo-EM grid preparation. Sample was concentrated to ~8mg/mL using 100kDa MWCO concentrators. Protein (3ul) was applied to plasma cleaned UltrAuFoil 1.2/1.3 300 mesh grids, blotted for 6s with 0 blot force, and plunge frozen into nitrogen cooled liquid ethane using a Vitrobot Mark IV (ThermoFischer).

### Cryo-EM data collection

Images were collected at 200 kV on a Talos Arctica electron microscope (ThermoFischer) with a K2 direct electron detector (Gatan) at a nominal pixel size of 1.15Å. Leginon software was used to automatically collect micrographs (*23*). The total accumulated dose was ~55 e-/Å^2^ and the defocus range was 0.8-1.5 um. Movies were aligned and dose-weighted using MotionCor2 (*24*).

### Image processing

Images were assessed for quality and edges of gold holes were masked using EMHP (*25*). CTF values were estimated using Gctf (*26*). Template-based particle picking was completed using FindEM template correlator (*27*). Particles were extracted using Relion 2.1 (*28*) then subjected to 2D classification using cryoSPARC (*29*). 130,054 particles corresponding to good class averages were selected for further data processing. An ab initio initial model was created in cryoSPARC followed by iterative angular reconstitution and reconstruction. The resulting density map was used as a seed for refinement of the data set in Relion 2.1. Resolution of the resulting map was 4.6Å. The map showed significant disorder in the LLR regions; however the map reveals that LRR regions arrange pairwise around a three-fold symmetry axis. As the transmembrane and extracellular domains were well-resolved, refinement was pursued imposing C3 symmetry and introducing a mask that excluded density outside of the well-defined, three-fold symmetric transmembrane/extracellular domains. Resolution of the resulting map was 4.0Å; transmembrane/extracellular domains were well-resolved whereas LRR regions were largely disordered. This map was then used to create suitable projections that were subtracted from particles, thereby creating a particle data set corresponding mostly to LRR densities. This new data set was then subjected to 3D classification in Relion 2.1 (K-means split of 12). One of the resulting classes showed order in the pairwise LRR arrangement around the three-fold symmetry axis. Particles corresponding to this class (25,719) were then refined locally around the previously obtained coordinate assignment imposing three-fold symmetry resulting in an LRR density map at 5.0Å resolution. Additionally ‒ due to the overall higher degree of order ‒ original particles corresponding to the 25,719 density-subtracted particles were refined under three-fold symmetry constraints. Resolution of the resulting map was 4.4Å.

### Model building and refinement

An initial model of an N-terminal portion of LRRC8A was generated with RobettaCM using innexin-6 (5H1Q) as a template structure (*11*, *30*). The LRRC8A topology was predicted using OCTOPUS (*31*). Predicted transmembrane regions were manually aligned to the transmembrane helices of the template structure 5H1Q (*11*). Intervening regions of LRRC8A were aligned to 5H1Q using BLASTp. 10,000 independent homology models were generated with RosettaCM and clustered using Calibur (*32*). The resulting model with the lowest Rosetta energy from the largest cluster was used as a guide for *ab initio* building of the transmembrane helices, extracellular domains, and intracellular linker domain. Sequence register was aided by bulky side chains and disulfide bonds in the extracellular domain. A Robetta-generated model of the LRRC8A LRR domain was docked into the EM density corresponding to the LRR of the outer subunit, which was better resolved than the inner subunit (*33*). This LRR model was adjusted manually to fit the density, then copied and docked into the LRR density of the inner subunit, followed by further adjustments. During the building process, manual building in COOT (*34*) was iterated with real space refinement using Phenix (*35*) or RosettaRelax (*36*). Structures were evaluated using EMRinger (*37*) and MolProbity (*38*). The final model contains residues 15-68, 94-174, 232-802 in the inner subunit and 15-68, 94-175, 214-802 in the outer subunit. Side chains of residues 15-21, 359-364, 787-802 of both subunits and 214-233 of the outer subunit were trimmed to Cβ because of limited resolution and lack of well-defined secondary structures in these regions. Structure figures were made in Pymol (*39*) and UCSF Chimera (*40*). Pore radii were calculated using HOLE (*41*). The APBS plugin in pymol was used to calculate surface representations of electrostatic potentials.

### Electrophysiology and cell culture

Electrophysiology experiments were completed with either Hela (ABCDE)^−/−^ cells or ShA KD Hela cells, described in Qiu et al., 2014 (*8*). ShA KD Hela cells cells were transfected with Lipofectamine 2000 and tested 1-4 days later (*8*, *9*). HeLa (ABCDE)^−/−^ cells were transfected 2-3 days earlier with SWELL1 constructs together with LRRC8C-ires-GFP in a 2:1 ratio (0.8 and 0.4 γ/ml for each coverslip). VRAC currents using a 2:1 ratio of SWELL1:LRRC8C were at least twice as large as those using a 1:1 ratio (data not shown). Only one cell per coverslip was tested for its response to hypotonic solution. In experiments on SWELL1 only and Y127C current, the extracellular solution contained (in mM) 90 NaCl, 2 KCl, 1 MgCl_2_, 1 CaCl_2_, 10 HEPES, 110 mannitol (isotonic, 300 mOsm/kg) or 30 mannitol (hypotonic, 230m0sm/kg), pH 7.4 with NaOH. Recording pipettes were filled with intracellular solution containing (in mM): 133 CsCl, 5 EGTA, 2 CaCl_2_, 1 MgCl^2^, 10 HEPES, 4 Mg-ATP, 0.5 Na-GTP (pH 7.3 with CsOH; 106 nM free Ca^2+^) and had resistances of 2-3 MΩ. Experiments testing R103F and T5 mutants used extracellular solutions described in Qiu et al., 2014 (*8*) (“bianionic”) and intracellular solution used in Syeda et al., 2016 (130mM CsCl, 10 HEPES, 4 Mg-ATP, pH 7.3). These were used to determine relative permeability P_I_/P_Cl_. An agar bridge was used between the ground electrode and the bath in all experiments.

---

**Fig. S1.**
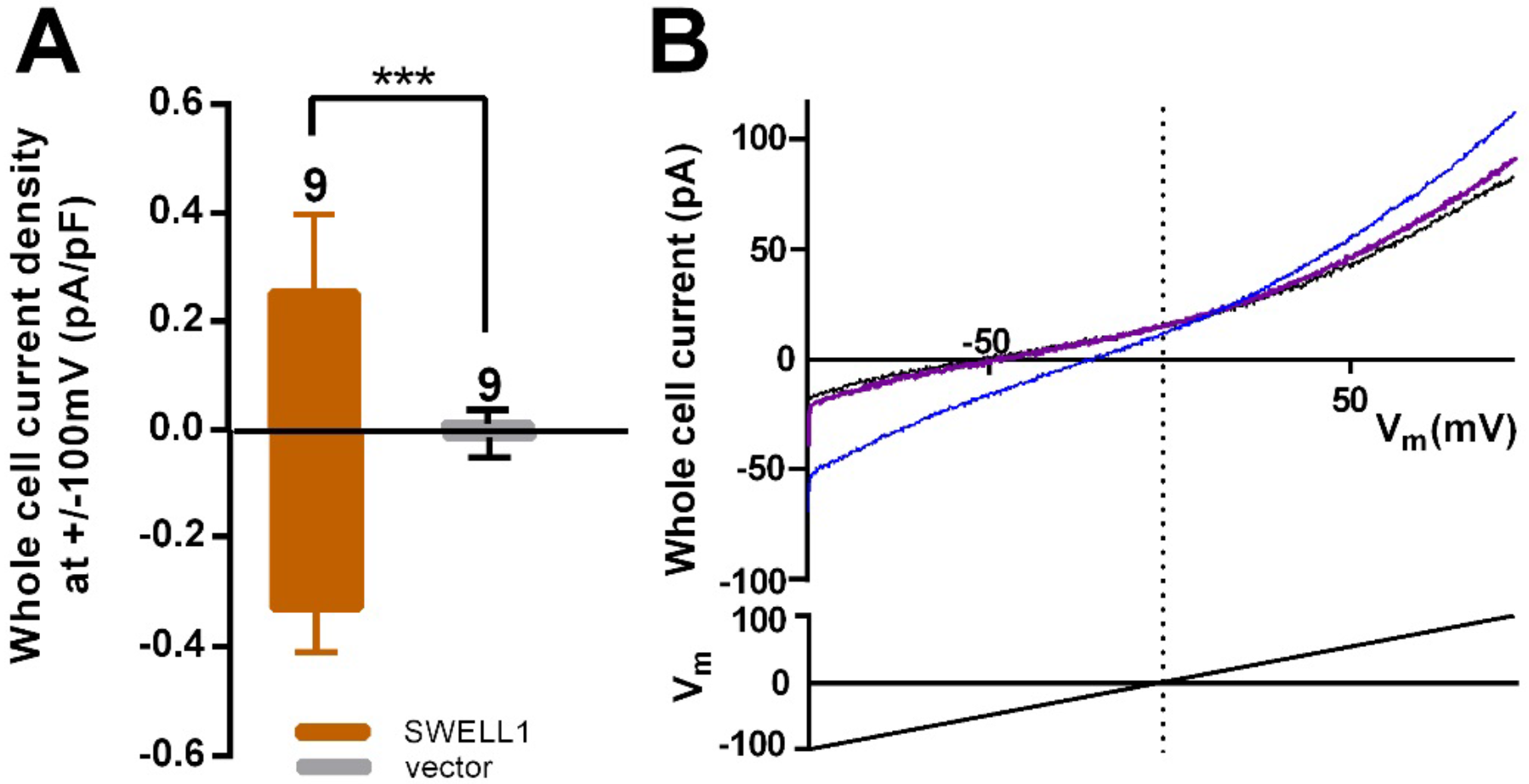
SWELL1 overexpression in cells lacking other LRRC8 subunits produces DCPIB-sensitive swelling-induced whole cell currents. (**A**) Hela (ABCDE)^−/−^ cells were transfected with SWELL 1-pIres-GFP or empty plres-GFP vector and tested for hypotonic (230 mOsm/kg) solution-induced currents using voltage ramp protocols (*8*). Whole cell current densities were determined at -100 and +100 mV before and 5-7min after hypotonic challenge. Shown are means ± S.E.M. for the number of cells indicated above the error bar. (**B**) Hypotonic solution-induced currents (blue) were blocked by DCPIB (20μM) (purple). Black trace is control. Traces were filtered at 1900Hz and averaged (ten sweeps each). Extracellular solution (in mM): 90 NaCl, 2 KCl, 1 MgCl2, 1 CaCl2, 10 HEPES, 110 mannitol (isotonic, 300 mOsm/kg) or 30 mannitol (hypotonic, 230mOsm/kg), pH 7.4 with NaOH. Recording pipettes were filled with intracellular solution containing (in mM): 133 CsCl, 5 EGTA, 2 CaCl2, 1 MgCl2, 10 HEPES, 4 Mg-ATP, 0.5 Na-GTP (pH 7.3 with CsOH; 106 nM free Ca2+) and had resistances of 2-3 MΩ.

**Fig. S2.**
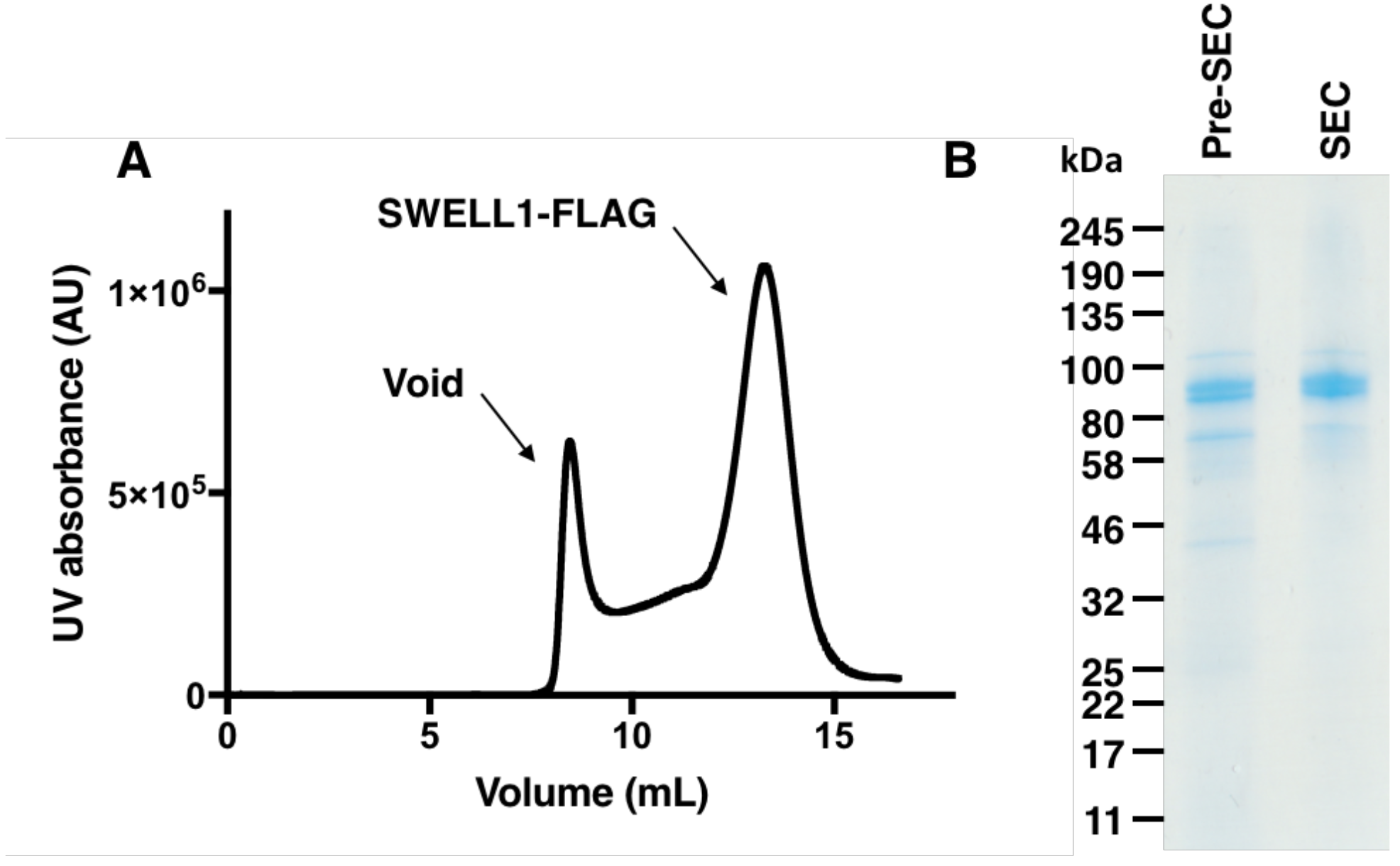
Purification of SWELL1-FLAG. (**A**) Gel filtration trace of SWELL1-FLAG after affinity purification. (**B**) SDS-PAGE gel of SWELL1-FLAG. Lane 1 is eluate after affinity purification (Pre-SEC) and lane 2 is combined fractions from SWELL1-FLAG peak post size exclusion. Expected molecular weight is 94 kDa.

**Fig. S3.**
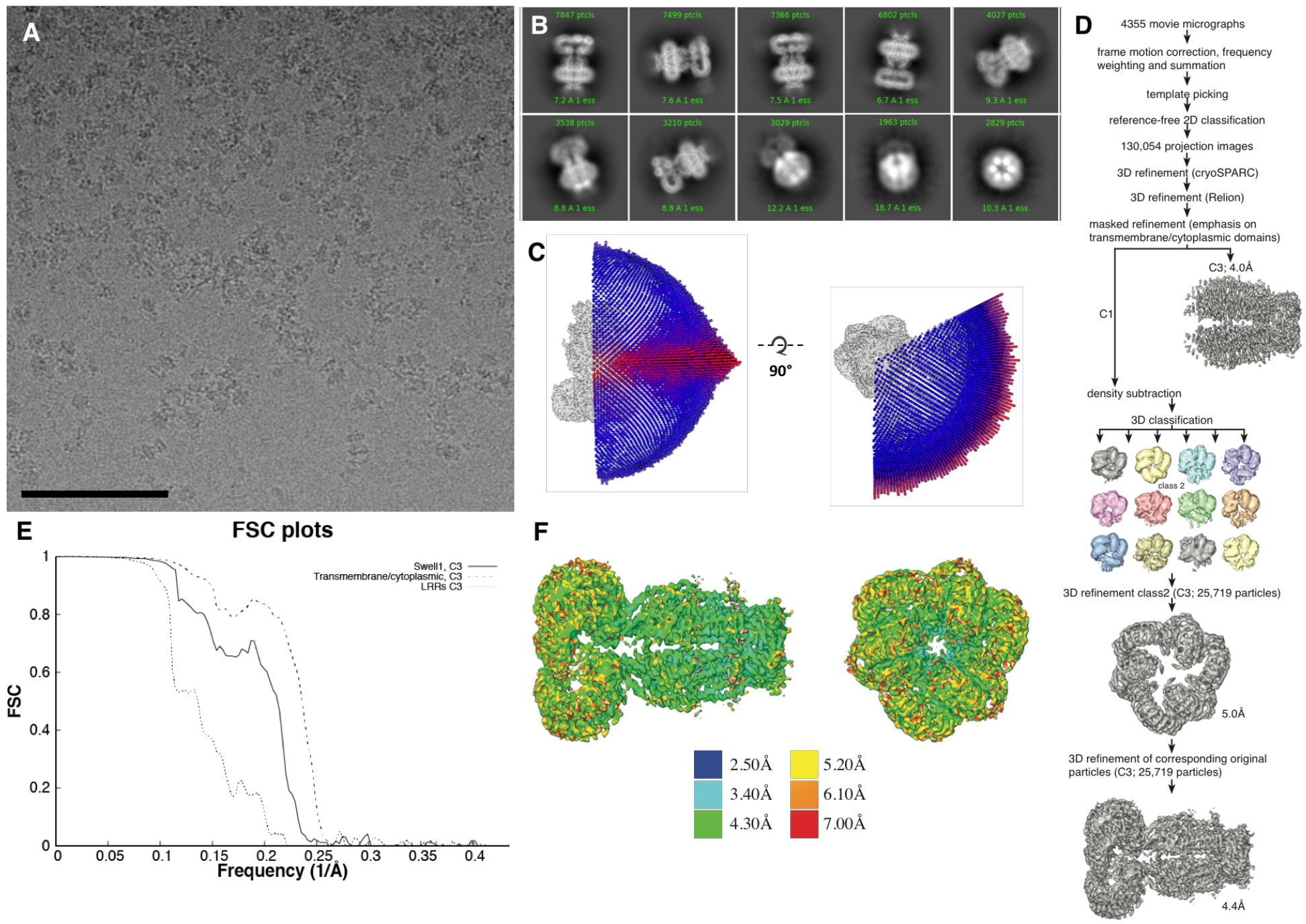
Cryo-EM data collection. (**A**) Representative aligned micrograph (scale bar, 100 μm). 4355 movies were collected of SWELL1-FLAG in vitreous ice. (**B**) Representative 2D classes showing range of orientations of SWELL1-FLAG particles. (**C**) Euler distribution for final map. Applied C3 symmetry was used, thus only one third of the sphere is shown. (**D**) Data-processing flow chart. (**E**) Fourier shell correlation between to independently refined cryo-data half sets (**F**) Local resolution estimates of the final reconstruction.

**Fig. S4.**
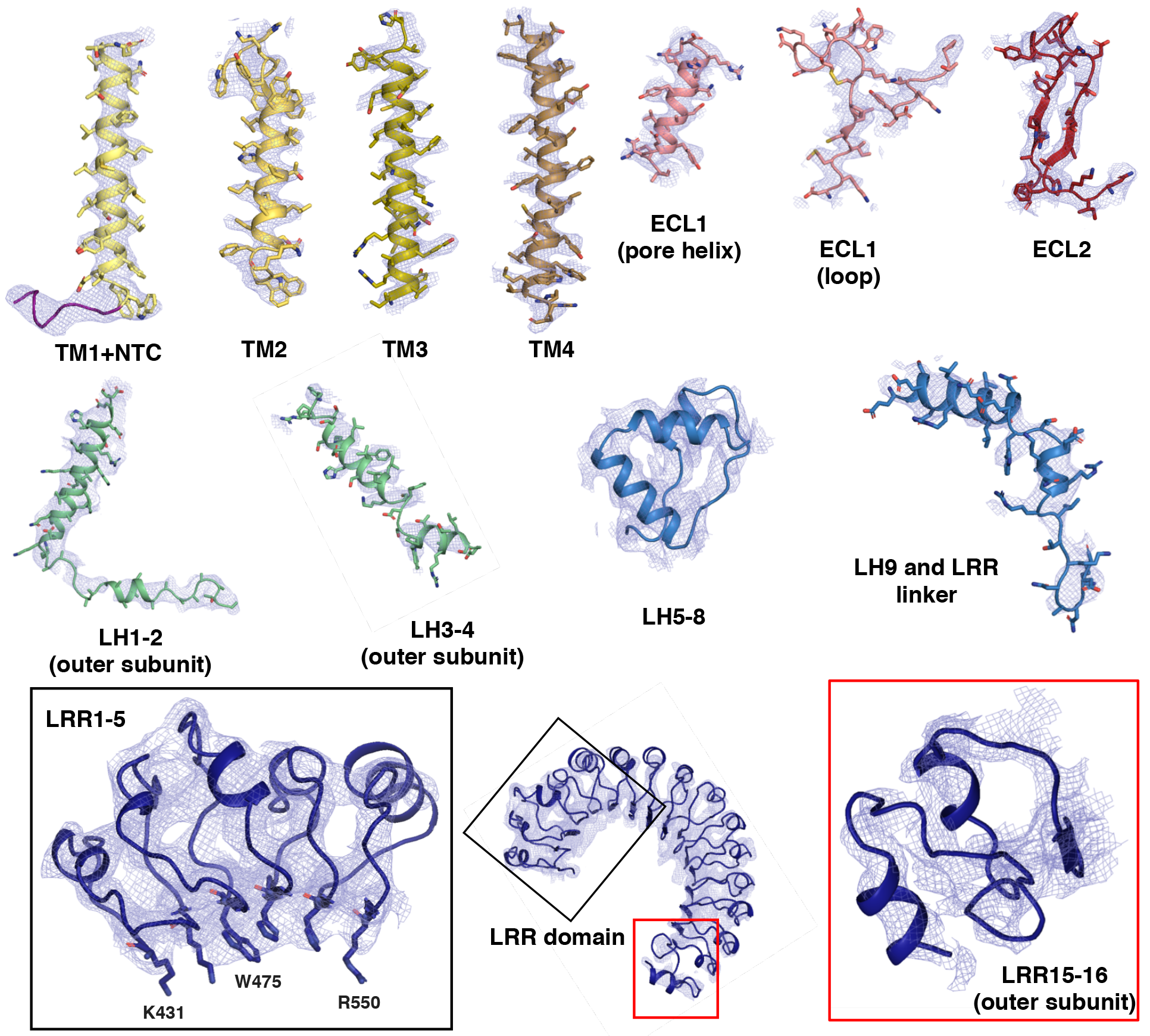
Model-to-map fit of electron density. Selected regions of the model are shown with superimposed electron density. Density is derived from the final C3 symmetry imposed map sharpened with a b-factor of-110 Å^2^. The cytosolic loop helices (LH1-4) and LRR 15-16 are from the outer subunit.

**Fig. S5.**
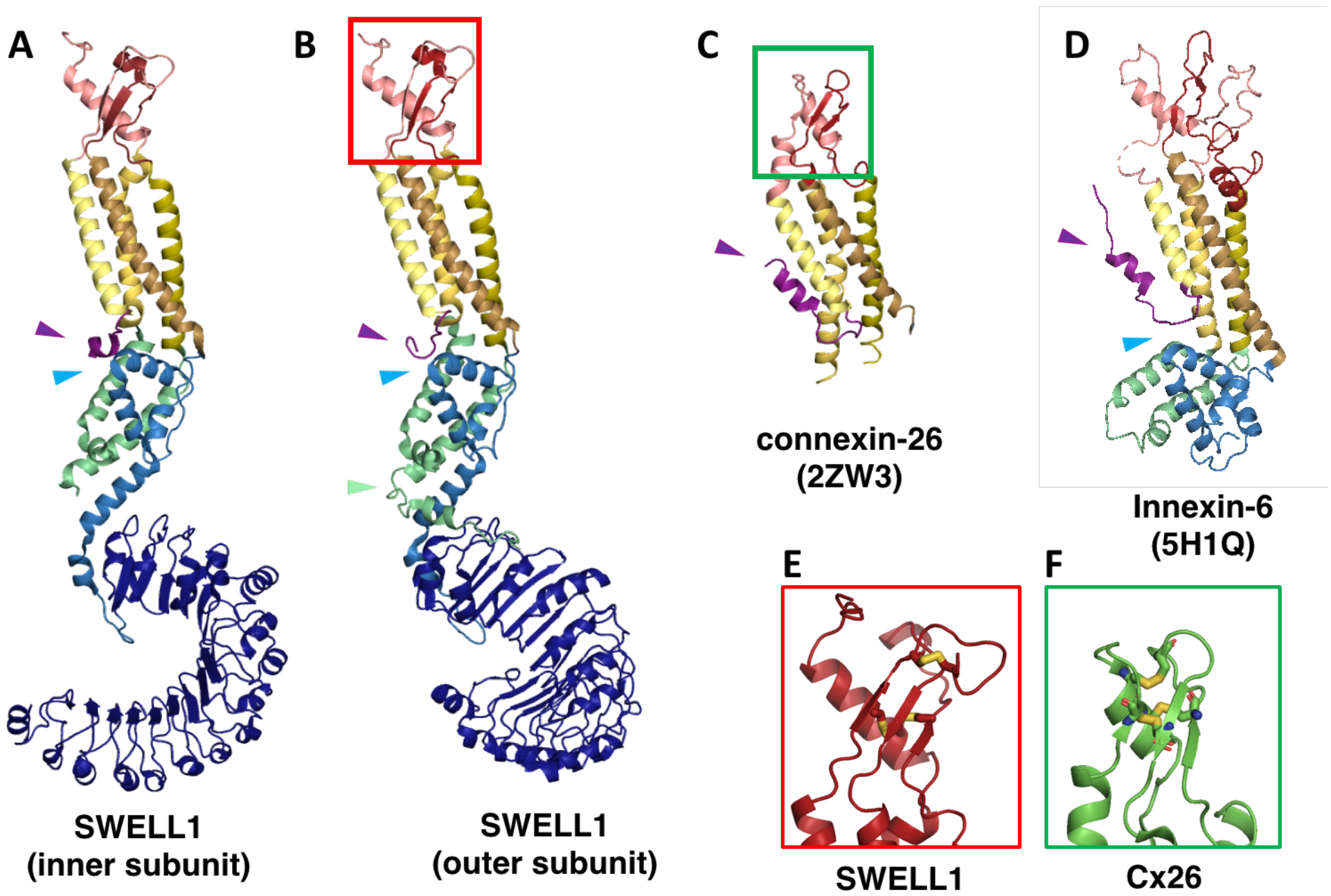
SWELL1 subunit pair and structural homology of SWELL1 structure to connexin-26 and innexin-6 structures. (**A-B**) Comparison of the SWELL1 “inner subunit” and “outer subunit” in a dimer pair. In the inner subunit, the TM2-TM3 linker contributes two long parallel helices (LH1 and LH4) with a flexible loop of res175-231 between them, while the outer subunit has an additional pair of parallel helices (LH2 and LH3) kinked to place them between the two subunits, though still with an unresolved loop of res176-213 between them. LH2, corresponding to residues R214-E236, sits atop of the LRR in the outer subunit (Green arrow). Additionally, there is a rotation of the LRR domains relative to the TM domains that allows them to dimerize. (**A-D**) The LRRC8 family has been shown to be related by weak sequence homology to pannexins (*42*), which are in turn related by structural homology to connexins and innexins. The transmembrane helices (TM1-4) of SWELL1 share the same order and arrangement as those of connexin-26 (Cx26; PDB: 2ZW3) (*10*) and innexin-6 (PDB: 5H1Q) (*11*) with TM1 closest to the central axis and TM3 and TM4 facing the membrane environment. An N-terminal helix in both Cx26 and innexin-6 creates a pore funnel that forms the narrowest constriction of the channel (*10*, *11*), which may be recapitulated in SWELL1 (Purple arrows) (**E-F**) The extracellular loops of SWELL1 (red box) share 3 structurally conserved disulfide bonds with connexin-26 (green box), as well as a three-strand antiparallel beta sheet composed of the antiparallel beta hairpin of ECL2, and a beta strand from ECL1 (*10*). (**A-B, D**) In SWELL1, cytoplasmic core formed by the intracellular linker domains share the same topology with portions of the C-terminal domains of distantly related innexin-6 (Blue arrows) (*11*).

**Fig. S6.**
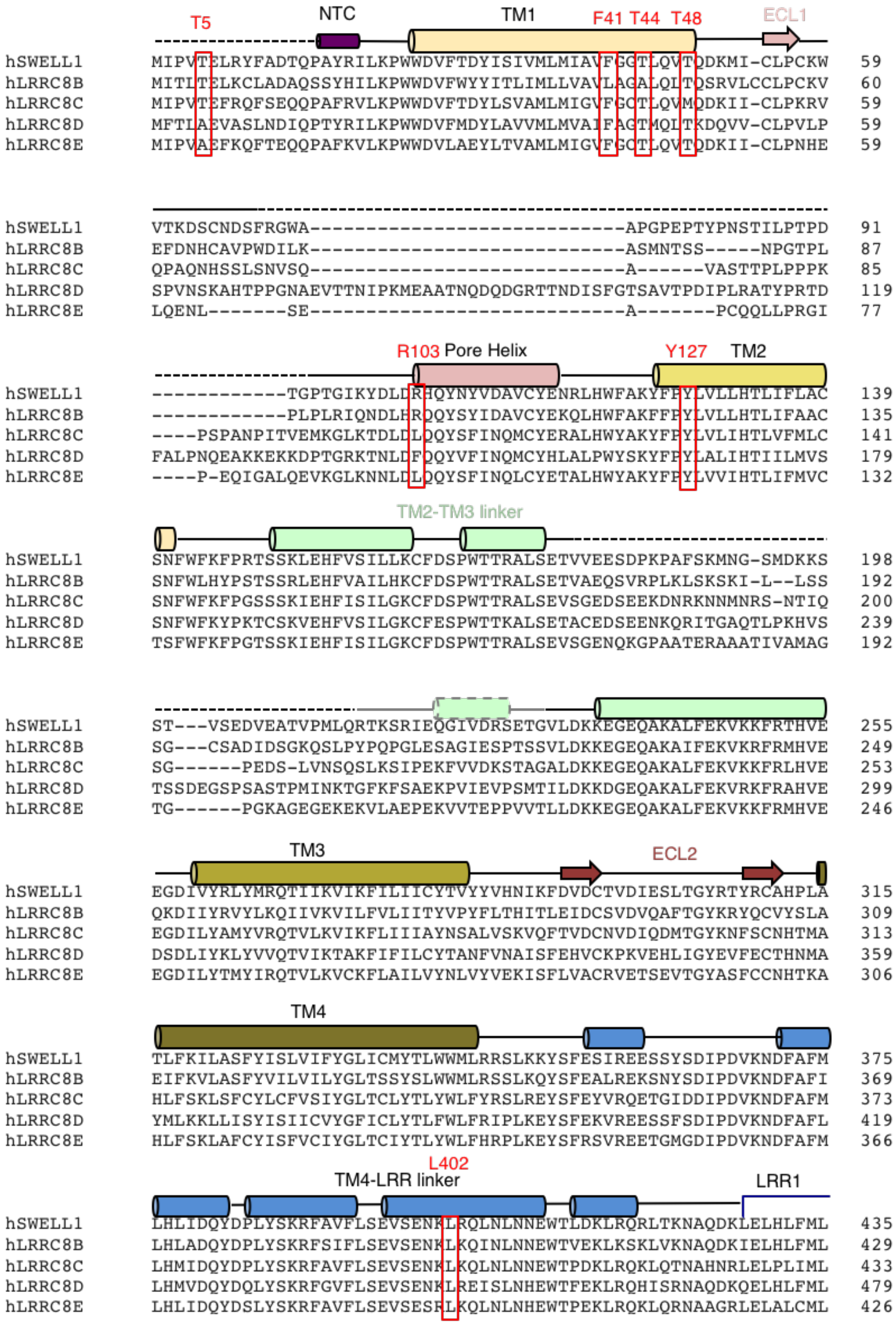
Alignment of LRRC8 subunits. Human LRRC8 family aligned using ClustalW. Secondary structure and domain assignments are annotated above. Residues mentioned in text are boxed red.

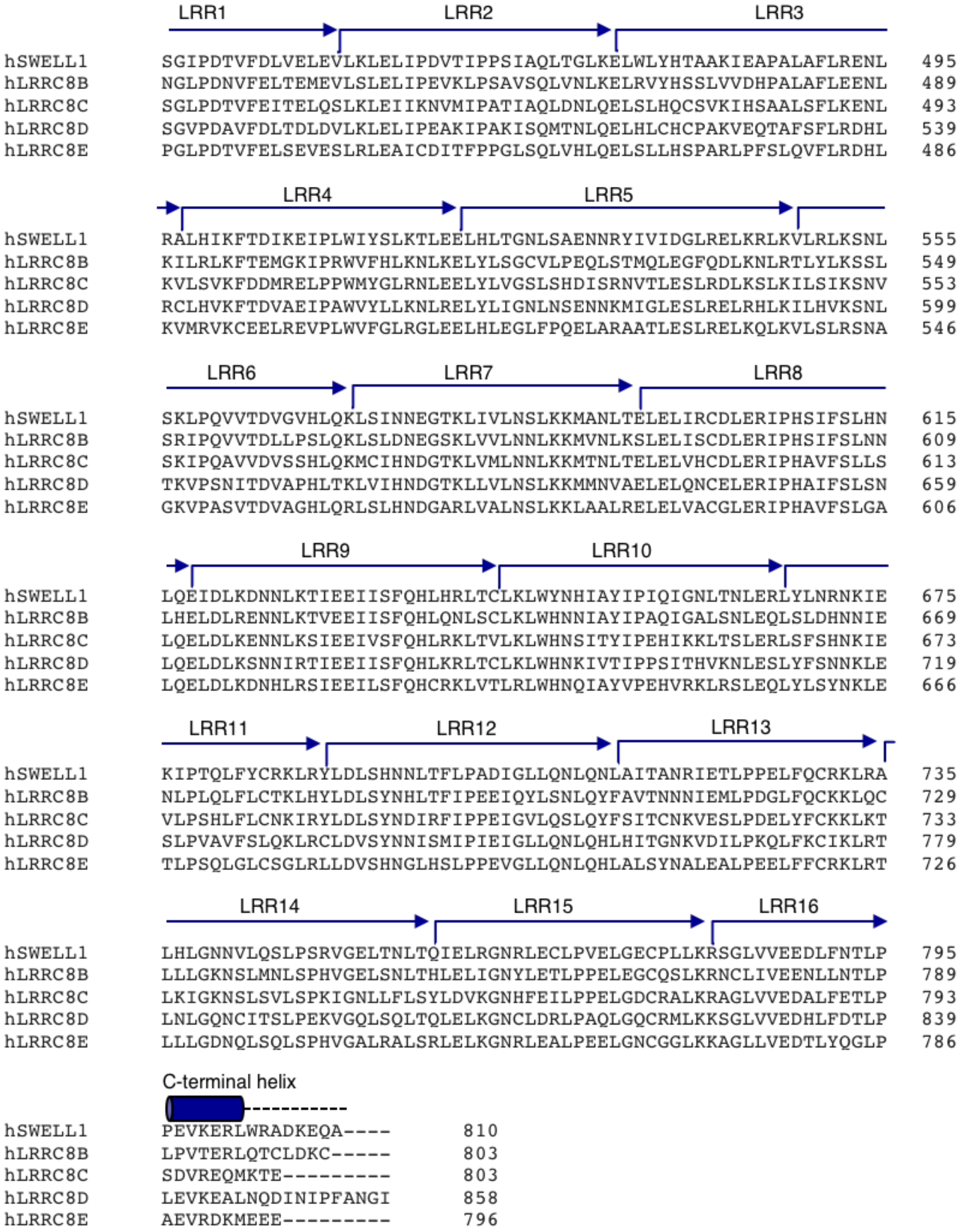

**Fig S7.**
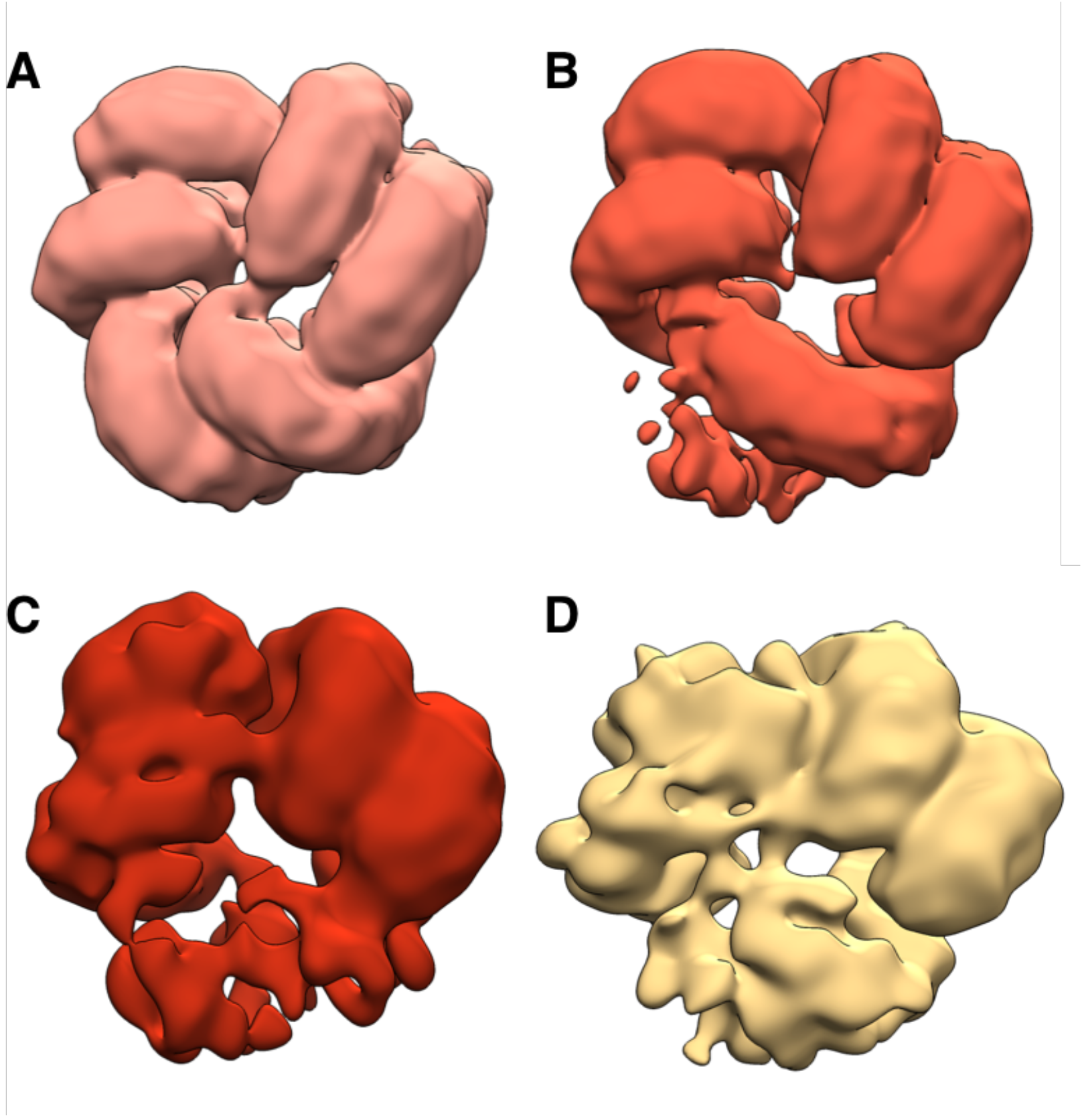
Flexibility in LRR domains observed during 3D classification. Bottom view of arrangement of LRR domains in representative subclasses of density-subtracted 3D classification (See methods). Precleaned particle picks were distributed into classes with clear organization of three (**A**; 25,607 particles), two and one half (**B**; 29,638 particles), two (**C**; 7,197 particles), and one (**D**; 8,071 particles) pairs of LRRs are observed. The highest resolution map was produced with the particles in the classes in which all three pairs are resolved, while the classes in which one or more LRR pairs are flexible produce lower resolution refinements.

**Fig S8.**
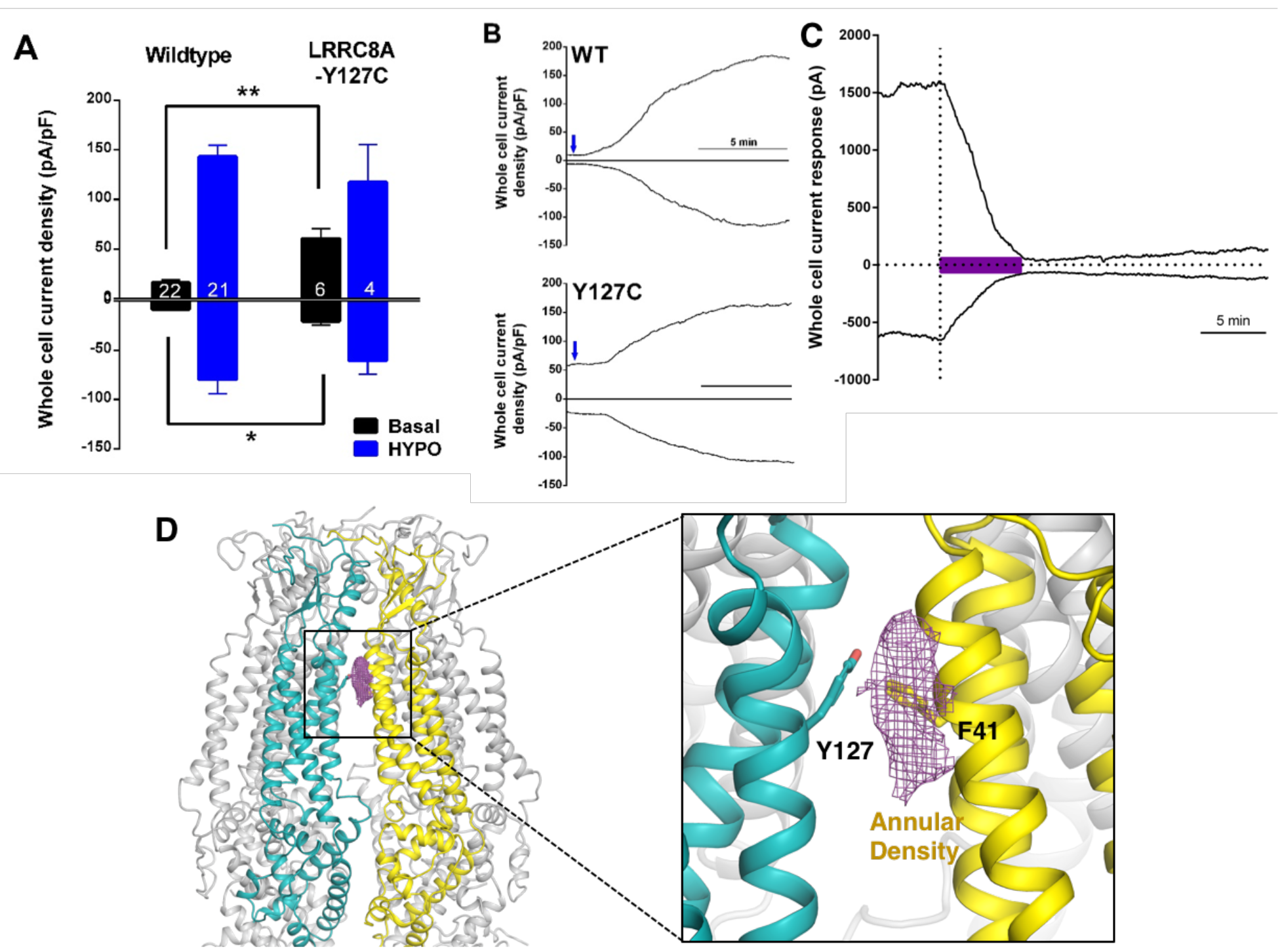
Role of interactions between subunits at the extracellular/membrane interface. (**A-C**) Large DCPIB-sensitive SWELL1-Y127C dependent currents were observed in the whole cell configuration within 10 sec of patch rupture. HeLa cells overexpressing SWELL1-wildtype or SWELL1-Y127C were challenged with voltage ramp protocol and after ~2 min exposed to hypotonic solution (230 mOsm/kg). (**A**) Basal currents were measured at -100 and +100 mV within 10 sec after patch rupture (black bars). The maximum current observed at -100 and +100 mV in the presence of hypotonic solution was determined (blue bars). Although VRAC expression was similar between WT and Y127C expressing cells, there was a significant increase in the basal current observed after patch rupture for Y127C-mediated I_Cl,swell_ compared to wildtype-mediated I_Cl,swell_ (p<0.01 at +100 mV; p< 0.05 at -100 mV; Student’s *t*-test). Shown are the means ± S.E.M. for the number of cells indicated. The combined basal and hypotonic solution-induced currents were similar (at +100 mV: 160 ± 12 pA/pF (wildtype) vs. 179 ± 39 pA/pF (Y127C); at -100 mV: -87 ± 14 pA/pF (wildtype) vs. -82 ± 14 pA/pF (Y127C); uncertainty in sums was used to calculate S.E.M.). Cm (cell capacitance) was used to calculate current density and determined as described in Qiu et al., 2014 (*5*). Cm did not differ between data sets (wildtype: 20.2 ± 0.7 pF (n=22); Y127C: 18.8 ± 1.7 pF (n=6); mean ± S.E.M.). V_rev_ for hypotonic solution-induced response was similar (wildtype: +7.3 ± 0.3 mV (n=21); Y127C: +6.7 ± 1.5 mV (n=4)). ShA KD HeLa cells were transfected with Lipofectamine 2000 and tested 1 day later as described (*8*). (**B**) Representative data from ShA KD HeLa cells transfected 1 day earlier with wildtype SWELL1 (top) and SWELL1-Y127C (bottom). Current density is plotted at +100 mV (top trace) and -100 mV (bottom trace). Only current densities from 30 sec prior to continuous hypotonic challenge (initiated at blue arrow) are shown but the basal current densities after rupture and before hypotonic challenge are similar. Extracellular solution contained (in mM) 90 NaCl, 2 KCl, 1 MgCl_2_, 1 CaCl_2_, 10 HEPES, 110 mannitol (isotonic, 300 mOsm/kg) or 30 mannitol (hypotonic, 230mOsm/kg), pH 7.4 with NaOH. Recording pipettes were filled with intracellular solution containing (in mM): 133 CsCl, 5 EGTA, 2 CaCl_2_, 1 MgCl_2_, 10 HEPES, 4 Mg-ATP, 0.5 Na-GTP (pH 7.3 with CsOH; 106 nM free Ca^2+^) and had resistances of 2-3 MΩ. (**C**) DCPIB (20uM) blocks basal whole cell currents in a ShA KD HeLa cell overexpressing SWELL1-Y127C. A similar result was observed in another cell. (**D**) The interface between two adjacent helix bundles of neighboring subunits near the extracellular side of the membrane. (Inset) F41 on TM1 and conserved Y127 on TM2 of the neighboring subunit form an interface. Also in this region, a putative lipid or detergent molecule sits between the helix bundles of neighboring protomers.

**Fig S9.**
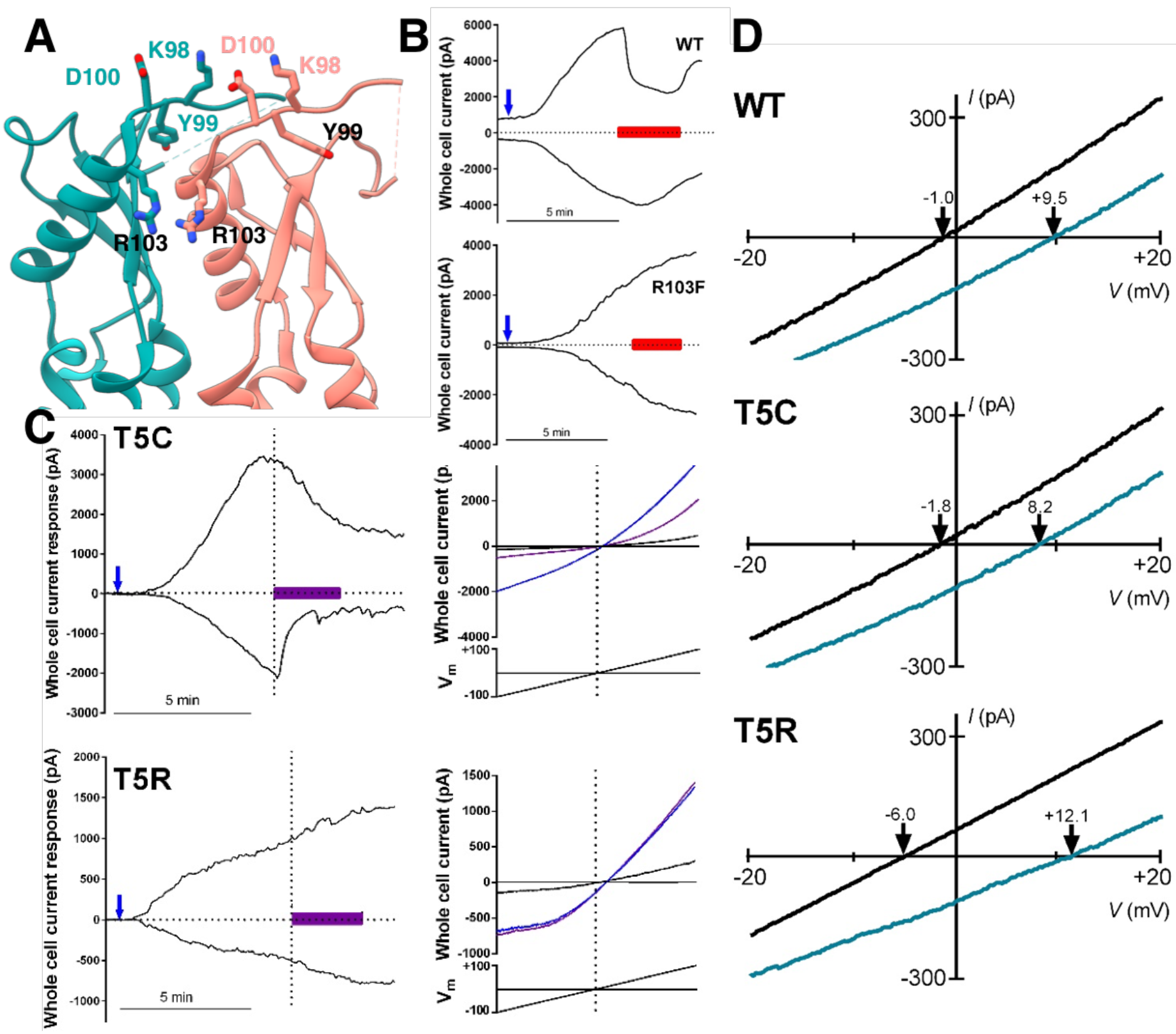
Additional characterization of selectivity and other pore properties. (**A**) Detailed view of neighboring extracellular loops with KYD motif and pore-constricting residue R103 labeled (**B**) Representative data of ATP block from HeLa (ABCDE)^−/−^ cells transfected 2-3 days earlier with wildtype SWELL1 (top) and SWELL1-R103F (bottom) together with LRRC8C in a 2:1 ratio. Currents at +100 mV (upper trace) and −100 mV (lower trace) are plotted. Blue arrow indicates addition of hypotonic solution; red bar, Na_2_ATP. Chloride solutions described in bianionic experiments (*8*, *9*) were used (Cl- in: 130mM; out: 88mM). (**C**) The polar MTS reagent MTSES applied extracellularly blocks SWELL1-T5C containing channels (top) but has no effect on T5R containing channels that are not modifiable by MTS reagents. Whole cell currents induced by hypotonic solution (230 mOsm/kg; blue arrows) in HeLa (ABCDE)^−/−^ cells heterologously expressing SWELL1-T5C (top) or SWELL1-T5R (bottom) with LRRC8C in a 2:1 ratio (0.8 and 0.4 γ/ml; see Qiu et al., 2014 (*8*)). MTSES (purple bar; 3.33mM) strongly reduced T5C-but not T5R-mediated currents in a manner consistent with covalent modification since reversibility of the T5C block was not observed during washout. Representative currents elicited by voltage ramps from −100 mV to +100 mV are shown (black, before; blue, in hypotonic solution before MTSES; purple, during MTSES). (**D**) Representative leak subtracted whole cell ramp-induced currents elicited by hypotonic solutions (230 mOsm/kg) containing either 88mM NaCl/10mM HEPES (green traces) or 88mM Na-Iodide/10mM HEPES (black traces) from HeLa (ABCDE)^−/−^ cells overexpressing WT, T5C or T5R mutants together with LRRC8C at 2:1 ratio. Bianionic solutions described in Qiu et al., 2014 (*8*) were used to determine relative permeability (P_I_/P_Cl_).

**Table S1.** Cryo-EM data collection, refinement and validation statistics.

|  |  |
| --- | --- |
|  | hSWELL1<br>(EMDB-XXXX)<br>(PDB ID XXXX) |
| **Data Collection and Processing** |  |
| Electron microscope | Talos Arctica |
| Electron detector | K2 Summit |
| Magnification | 36,000 |
| Voltage (kV) | 200 |
| Electron exposure (e-/Å<sup>2</sup>) | 5.9 |
| Defocus range (µm) | 0.8-1.5 |
| Pixel Size (Å) | 1.15 |
| Symmetry imposed | C3 |
| Initial particle images (no.) | 316,206 |
| Final particle images (no.) | 25,719 |
| Map resolution (Å) | 4.4 |
| FSC threshold | 0.143 |
| Map resolution range (Å) | 3.5 - >7.0 |
| Map sharpening *B* factor (Å<sup>2</sup>) | -110 |
| **Model building and refinement** |  |
| Initial models used | RobettaCM generated model using<br>PDB ID 5H1Q as template structure |
| Model composition |  |
| Protein Chains | 6 |
| Protein Residues | 4273 |
| Ligands | 0 |
| R.m.s. deviations |  |
| Bond Lengths (Å) | 0.01 |
| Bond angles (°) | 1.18 |
| Ramachandran plot |  |
| Favored (%) | 85.01 |
| Disallowed (%) | 0.16 |
| Validation |  |
| MolProbity score | 2.02 |
| Clashscore | 6.57 |
| Poor rotamers (%) | 0.08% |
| EMRinger score | 1.1 |

